# Vascular-water-exchange MRI (VEXI) enables the detection of subtle BBB breakdown in Alzheimer’s disease without MRI contrast agent

**DOI:** 10.1101/2022.09.27.509514

**Authors:** Yifan Zhang, Yue Wang, Zhaoqing Li, Zejun Wang, Juange Cheng, Xiaoyan Bai, Yi-Cheng Hsu, Yi Sun, Shiping Li, Jiong Shi, Binbin Sui, Ruiliang Bai

## Abstract

Blood-brain barrier (BBB) impairment is an important pathophysiological process in Alzheimer’s disease (AD) and a potential biomarker for early diagnosis of AD. However, most current neuroimaging methods assessing BBB function need the injection of exogenous contrast agents (or tracers), which limits the application of these methods in a large population. In this study, we aim to explore the feasibility of vascular water exchange MRI (VEXI), a diffusion-MRI-based method to assess the BBB permeability to water molecules without using a contrast agent, in the detection of the BBB breakdown in AD. We tested VEXI on a 3T MRI scanner on three groups: AD patients (AD group), mild cognitive impairment (MCI) patients due to AD (MCI group), and the age-matched normal cognition subjects (NC group). Interestingly, we find that VEXI can detect the BBB permeability to water molecules increase in MCI and this BBB breakdown happens specifically in the hippocampus. This BBB breakdown gets worse and extends to more brain regions (orbital frontal cortex and thalamus) from MCI group to the AD group. Furthermore, we find that the BBB breakdown of these three regions detected by VEXI is correlated significantly with impairment of respective cognitive domains independent of age, sex and education. These results suggest VEXI is a promising method to assess the BBB breakdown in AD.

**Highlights:**

- The vascular water exchange MRI (VEXI) is a contrast-agent-free method to assess BBB permeability
- BBB breakdown happens specifically in the hippocampus, orbital frontal cortex, and thalamus in AD
- BBB breakdown detected by VEXI is significantly correlated with cognitive dysfunction

## 1. Introduction

Increasing evidence has shown that blood-brain barrier (BBB) impairment is a contributing factor in the pathophysiology of Alzheimer’s disease (AD) (Iadecola, 2017; Sweeney et al., 2018). BBB is a continuous endothelial membrane within brain microvessels that seals cell-to-cell contacts, regulates the delivery of important nutrients to the brain, and prevents neuro-toxins from entering the brain (Zlokovic, 2008). It also has a clearance function to remove excess substances from the brain. BBB structure and function can be assessed with neuropathology in postmodern samples (Arvanitakis et al., 2016; Toledo et al., 2013), neuroimaging methods (Kisler et al., 2017; Montagne et al., 2015; Sweeney et al., 2018; van de Haar et al., 2017, 2016), and cerebrospinal fluid biomarkers (Iturria-Medina et al., 2016; Montagne et al., 2015; Sagare et al., 2015; Sweeney et al., 2018), among which only neuroimaging methods could provide the spatial distribution features of BBB impairment *in vivo*. Studies using these neuroimaging methods, including positron emission tomography (PET) and MRI, have produced fruitful results of increased BBB permeability in different brain regions in AD patients (Minoshima et al., 1997; van de Haar et al., 2017, 2016; Wang et al., 2006) and its relationship with brain structural changes (e.g. atrophy) (Nation et al., 2019; Zhang et al., 2019). However, most conventionally used neuroimaging methods assessing BBB permeability in AD utilize the intravenous administration of contrast agents (or radioactive tracer). A non-invasive and contrast-agent-free neuroimaging method to characterize BBB permeability is still highly desirable.

Recently, there are several newly developed MRI methods aiming to measure the water exchange across BBB, which provides a novel means to assess BBB permeability without the usage of contrast agent (Dickie et al., 2020). These contrast-agent-free MRI techniques mainly include two different types of approaches. One is based on arterial spin labeling (ASL) technique, which uses ASL to label intravascular water and then monitor the dynamic change of labeled water in intravascular and extravascular spaces by using diffusion (He et al., 2018, 2012; Wang et al., 2007), multiple echo (Ohene et al., 2019), or phase contrast (Lin et al., 2018) methods. The second approach utilizes the filter-exchange imaging (FEXI), a technique adapted from diffusion exchange spectroscopy (DEXSY) for clinical applications (Lasič et al., 2011; Nilsson et al., 2013). By exploring the intravoxel incoherent motion (IVIM) of capillary water, designing a proper diffusion weighting to filter out intravascular water specifically and then quantitatively monitoring the water exchange between intra- and extravascular space via the second diffusion encoding, FEXI shows the capacity for measuring the water exchange across BBB in human (Bai et al., 2020). More encouragingly, a recent study shows that the water exchange across BBB is a more sensitive biomarker in the detection of subtle BBB breakdown than the conventional biomarker of contrast agent leakage from BBB in an AD rat model, as water molecule is much smaller than MRI contrast agents and could potentially be more sensitive to BBB leakage (Dickie et al., 2019). However, it is still unknown if it is feasible to use the water-exchange based MRI method to detect the BBB leakage in AD patients.

In this study, we aim to explore the feasibility of the contrast-agent-free MRI method in the detection of subtle BBB impairment in AD patients. For this purpose, the FEXI-based vascular-water-exchange MRI (VEXI) was implemented on a 3T clinical MRI scanner to assess the BBB permeability to water molecules. MRI and cognitive function assessments were performed on three groups: AD patients (AD group), mild cognitive impairment due to AD patients (MCI group), and the age-matched normal cognition subjects (NC group). Both MCI and AD groups were diagnosed based on clinical criteria and positive amyloid-beta (Aβ) deposition confirmed by PET.

## 2. Methods

### 2.1. Study Participants

This study was a sub-study of an ongoing prospective community-based cohort study of the China National Clinical Research Center Alzheimer’s Disease and Neurodegenerative Disorder Research (CANDOR). CANDOR was started in July 2019 and planned to enroll one thousand and five hundred participants, including individuals with NC, MCI, and dementia (including AD).

This sub-study recruited participants from March 2021 to January 2022. All the AD and MCI participants were recruited from Beijing Tiantan Hospital and the NC participants were recruited from the local communities. Demographic information, past medical history, social and family history were collected. All participants underwent detailed cognitive assessments, including the battery of neuropsychological tests such as Mini-Mental State Examination (MMSE), Montreal Cognitive Assessment (MoCA), Clinical Dementia Rating (CDR), Rey Auditory Verbal Learning Test (RAVLT) etc., and brain MRIs. All enrolled participants for this study were (1) subjectively normal in NC groups, or diagnosed as MCI due to AD or AD. The diagnosis of MCI due to AD and AD was based on the National Institute on Aging-Alzheimer’s Association guidelines for AD (Albert et al., 2011; McKhann et al., 2011) and positive amyloid status confirmed by ^11^C-labeled Pittsburgh Compound-B (PIB) PET-CT; (2) aged 40-100 years-old, (3) had at least 3 years of elementary-school education and could complete the neuropsychological tests independently; (4) had no conditions known to affect cognitive function, such as alcoholism, uncontrolled depression or other psychiatric disorders, Parkinson’s disease, epilepsy, stroke and etc.

### 2.2. Standard Protocol Approvals, Registrations, and Patient Consents

This protocol was approved by the Institutional Review Board of Beijing Tiantan Hospital (approval number: KY 2019-004-007) and was in accordance with relevant guidelines and regulations. Written informed consent was obtained from each participant.

### 2.3. Principles of vascular-water-exchange MRI (VEXI)

VEXI characterizes the BBB permeability to water molecules, which is illustrated in **Figure 1**. It is a specific type of diffusion-based FEXI adapted for measuring the water exchange across BBB (Bai et al., 2020). Briefly, it contains three blocks, including the filter block, mixing block, and detection block. The filter block is a pulsed gradient spin echo (PGSE) with diffusion weighting *b*_f_, in which *b*_f_ (= 250 s/mm^2^) is optimized to filter the intravascular water magnetization showing nearly one-fold larger apparent diffusivity than the extravascular water magnetization due to IVIM (Le Bihan, 2019). After the first filter block, the remaining magnetization in the transverse plane is stored back in the longitudinal direction and kept for a certain mixing time (*t*_m_), during which water molecules exchange across BBB (mixing block). Finally, the filtered and mixed magnetization is put back to the transverse plane with the third 90° pulse and the apparent diffusivity (*ADC*′) of the mixed water pools is measured in the second PGSE block (detection block). A pair of identical gradients is implemented before the second 90 pulse and after the third 90 pulse such that the stimulated echo is formed after the second gradient and the unwanted magnetization (e.g., the inflowing blood from neighboring slices) is eliminated.

**Figure 1.**
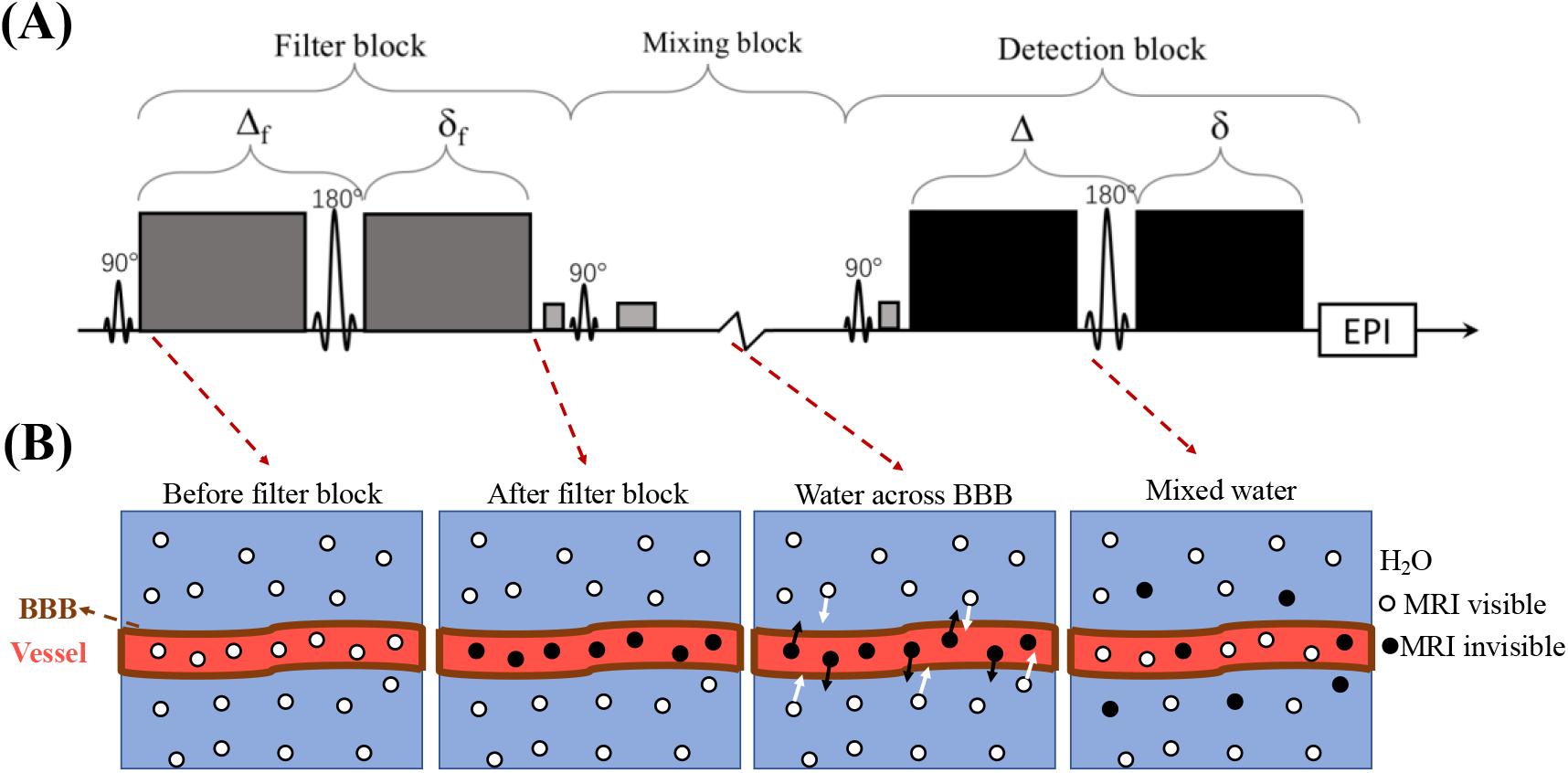
Overview of the principles of VEXI implemented in this study. (A) VEXI pulse sequence. (B) Illustration of the MRI signal evaluation of intravascular and extracellular water molecule during VEXI acquisition. The first pulsed gradient spin echo (PGSE) block with diffusion weighting *b*_f_ is served as the filter of intravascular water pools, which showing large apparent diffusivity due to IVIM. After the first filter block, the remaining magnetization in the transverse plane is stored back in the longitudinal direction (mixing block) and kept for a certain mixing time (*t*_m_), during which water molecules exchange across BBB. Finally, the filtered and mixed magnetization is put back to the transverse plane with the third 90° pulse and the apparent diffusivity (*ADC*′) of the mixed water pools is measured in the second PGSE block. In VEXI, *ADC*′ were acquired at several *t*_m_ (more water molecules exchange across BBB in longer *t*_m_) and the apparent water exchange rate across BBB (AXR_BBB_) could then be extracted from *ADC*′(*t*_m_) signal.

In VEXI, *ADC*′ were acquired at several *t*_m_ (more water molecule exchange across BBB in longer *t*_m_) and the apparent water exchange rate constant across BBB (AXR_BBB_) could then be extracted from *ADC*′(*t*_*m*_) signal (**Figure 1B**). Here it is assumed that the water molecules in brain tissue can be separated into “slow” (s, extravascular water pool) and “fast’” (f, intravascular water pool, IVIM) diffusion components with apparent diffusivities *D*_s_ and *D*_f_, and equilibrium fractional populations 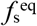 and 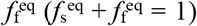, respectively. Water molecules are in exchange between the two water pools across BBB with a water exchange rate constant *k*_fs_ from the intravascular to the extravascular pool, and 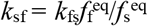 from the extravascular to the intravascular water pool. At a fixed *b*, the acquired MRI signal *S* is a function of the diffusion weighting in the detection block (*b*_d_) and *t*_m_ (Lasič et al., 2011; Nilsson et al., 2013),

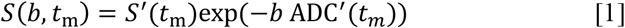

where *S*′(*t*_m_) takes into consideration the effects of longitudinal relaxation during *t*_m_ and the apparent diffusion coefficient *ADC*′(*t*_*m*_) is determined by the two water components’ apparent diffusivity and exchange rate, using an approximation that applies when *b* approaches zero,

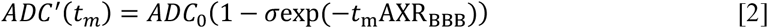

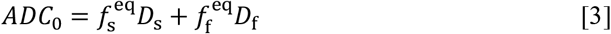

Here it is assumed that exchange is slow on the time scale of the duration of the PGSE blocks and that longitudinal relaxation rates are the same for both components. *σ* is the filter efficiency that quantifies the *ADC*′ reduction at *t*_m_ = 0. AXR_BBB_ is the apparent water exchange rate constant across BBB and its relation with the intravascular water efflux rate constant *k*_fs_ is,

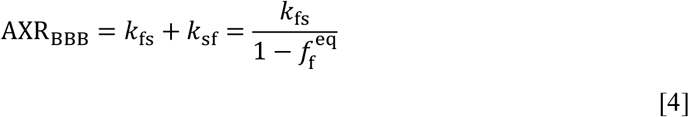

and AXR_BBB_ approaches to *k*_fs_ in gray or white matter as the intravascular water fraction 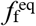 is generally small (e.g., < 5%) in healthy brain or the degenerated brain. In this study, AXR_BBB_ is taken as the quantitative parameter characterizing BBB permeability to water molecules.

### 2.4. MRI protocols

MRI scans were performed with a 3.0T MRI clinical scanner (MAGNETOM Prisma, Siemens Healthcare, Erlangen, Germany), using a Nova 64-channel head RF coil. MRI scans included 3D MPRAGE *T*_1_-weighted images, diffusion-weighted images and VEXI. The *T*_1_-weighted images were acquired with 1.0 × 1.0 × 1.0 mm^3^ resolution, TE/TR = 2.01/2000 ms, flip angle 8°, inversion time 880 ms. To characterize diffusion anisotropy, a diffusion tensor imaging (DTI) protocol was performed with TE/TR = 70/2800 ms, 5 repetitions on *b* = 0 s/mm^2^ and 50 directions with single repetition for each direction at *b* = 1000 s/mm^2^. The voxel size for the DTI images was 2.0 × 2.0 × 2.0 mm^3^.VEXI was performed at XZ direction with *b*_f_ = 250 s/mm^2^ and two *b* values in the detection block (*b*_d_ = 0 s/mm^2^ with 6 repetitions and 250 s/mm^2^ with 10 repetitions). Imaging was repeated with three mixing time (*t*_m_): 25, 200, and 400 ms. VEXI was also acquired with *b*_f_ = 0 s/mm^2^ and shortest *t*_m_ (25 ms), echo time in the filter block TE_f_ = 26 ms, echo time in the detection block TE = 37 ms, timings of the gradients in the filter block Δ_f_/δ_f_ = 11.5 ms / 6.3 ms, and timings of the gradients in the detection block Δ/δ = 14.4 ms / 8.6 ms. For the VEXI protocol, the voxel size was 3.0 × 3.0 × 5.0 mm^3^, and the number of slices was 20.

### 2.5. VEXI data processing

All DTI and VEXI data underwent pre-processing including motion and eddy current distortion correction in TORTOISE (Pierpaoli et al., 2010). For the DTI data, pre-processed DWIs were fit to non-linear DTI model and DTI metrics including mean diffusivity (MD) were generated with TORTOISE. For the VEXI data, the model fitting was performed with in-house programs developed in MATLAB (2018B, The MathWorks Inc., Natick, Massachusetts). Here, the ADC′(*t*_*m*_) values were computed from the measurements with two *b* values in the detection block (*b*_d1_ and *b*_d2_) at each *t*_m_, according to

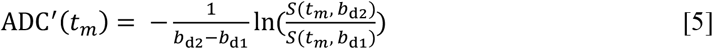

where *S*(*t*_*m*_, *b*_d1_)and *S*(*t*_*m*_, *b*_d2_) are the VEXI signals acquired at *b*_d1_ and *b*_d2_, respectively. Then the ADC′(*t*_*m*_) at three *t*_m_ and *b*_f_ = 0 (taken as equilibrium state, *i*.*e*., *t*_m_ = +Inf) were fitted to Eq. [2] with the trust-region nonlinear least-squares algorithm in each voxel to calculate the VEXI derived parameters, AXR_BBB,_ ADC and *σ*.

### 2.6 Region-of-interest (ROI) analysis

We then generated a study-specific template with the *T*_1_-weighted images of all subjects in NC group, using the ANTs script (Avants et al., 2010). DTI data with *b*=0 s/mm^2^ and VEXI data with *b*_f_=0 s/mm^2^ and *b*_d_=0 s/mm^2^ were linear registered to *T*_1_-weighted images using ANTs (Avants et al., 2011). Afterwards, the *T*_1_-weighted images in the native space were registered to our study-specific template using ANTs SyN algorithm. Parameter maps (AXR_BBB,_ ADC and *σ*) were then iteratively aligned to the template by applying the linear registration transformation matrix and the deformation field generated above. In addition, we nonlinearly registered the MNI152 *T*_1_-weighted template (Mazziotta et al., 2001, 1995) to our template to transform some atlas defined in MNI152 template into our study-specific template space.

Region-of-interest (ROI) masks were extracted from Brainnetome Atlas (Fan et al., 2016) and JHU white matter Atlas (Mori et al., 2005). As shown in **Figure 2**, a total of 5 grey matter ROIs including the hippocampus, caudate nucleus, thalamus, striatum, and orbital frontal cortex (OFC) and 3 white matter ROIs including subcortical watershed white matter fibers, corpus callosum and internal capsule were chosen according to other studies in AD or MCI (Montagne et al., 2015). All the ROIs were bilateral. Several steps were implemented to reduce the potential confound induced by brain atrophy, which includes (1) one experienced neurologist was invited to readjust and double-check the hippocampus and other brain ROIs based on neuroanatomy, (2) the 3D structural images (T1 weighted) were segmented into CSF, gray matter, and white matter, and these selected brain ROIs (like hippocampus) were readjusted to remove the CSF voxels based on the segmented results; (3) voxels with MD larger than 2.0 um^2^/ms were also considered as CSF voxels and further removed from these brain ROIs. The parametric metrics in each ROI were calculated as the median of all voxels in this ROI.

**Figure 2.**
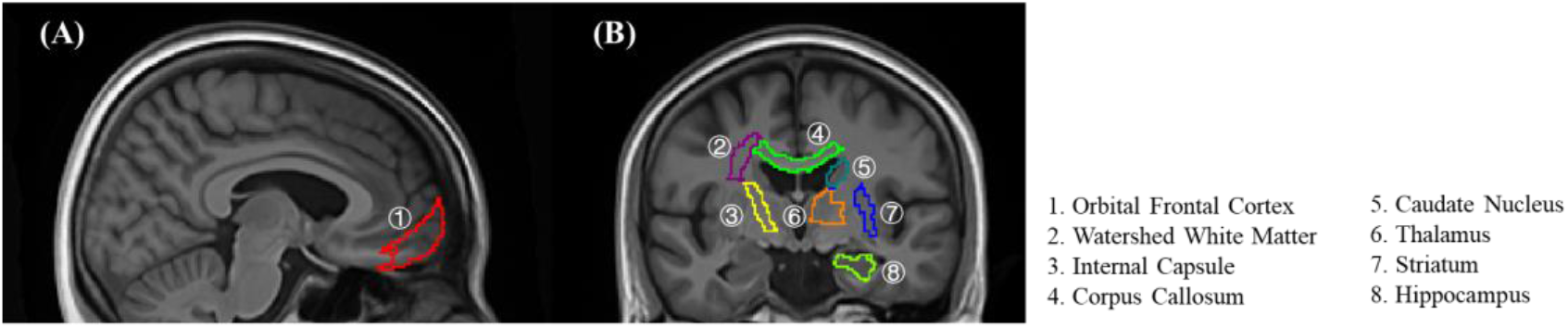
Eight regions of interest based on Brainnetome atlas and JHU white matter atlas in the customized-template space. Here different ROIs were represented with different numbers.

To double check the results obtained in the template space, the same ROI-level analysis was also performed in the subjects’ native space. The 8 selected brain regions were manually drawn by one experienced neurologist on each subject’s native space. CSF voxels were removed using the same methods in the template space. The parametric metrics in each ROI were also calculated as the median of all voxels in this ROI.

### 2.7 Volumetric analysis

For the regional level volumetric analysis, all the *T*_1_-weighted images were processed through the voxel-based-morphometry (VBM) pipeline in SPM12, which has been described previously (Ashburner and Friston, 2000). Briefly, we created our study-specific DARTEL template based on the template generated above. Then each *T*_1_-weighted image was segmented into GM, WM and CSF using the segmentation function in SPM12. Segmented images were then aligned to the DARTEL template iteratively, spatially normalized, modulated and smoothed with an 8mm Gaussian kernel. Volumes for the ROIs were calculated by summing all the white and grey matter voxels in the ROI from the pre-processed images. To correct for the head size, the total intracranial volume (TIV) was calculated as the sum of all voxels across the grey matter, white matter, and CSF segmented images. Then ROI volume was corrected by simply dividing by TIV, a widely used method for volumetric correction.

### 2.8 Statistics

Prior to performing statistical analyses, we first screened for outliers using the Grubbs’ test by applying a significant level of α=0.01 (Nation et al., 2019). All continuous variables were checked for normality through examination of skewness and kurtosis. Log10-transformations were applied where departures of normality were identified. Distribution normalization was confirmed before parametric analyses. Unpaired Students’ t-test was performed to compare the median AXR_BBB_ in each ROI between NC group and MCI group. One-way analysis of variance (ANOVA) was performed to explore the AXR_BBB_ change in each ROI among the three groups. One-way ANOVA was also employed to assess the difference in age, education levels and the scores of neurological tests, and the median ADC, *σ* and volume in all the ROIs among NC, MCI, and AD groups. Categorical variables were analyzed by Pearson’s χ^2^ tests. Correlation analysis between AXR_BBB_ and cognitive functions were performed with linear regressions, with age, sex and education level controlled.

All statistical analyses were performed in GraphPad Prism8.

## 3. Results

### 3.1 Subjects’ clinical characteristics

From March 2021 to January 2022, 52 participants were recruited for this sub-study from Beijing Tiantan Hospital and the local communities, including 27 healthy controls, 14 with MCI due to AD and 11 with AD. The baseline information of the subjects was showed in **Table 1**. There was no difference in age, gender and education level among groups. Statistical significance among groups was found in the neuropsychological tests, including MMSE, MoCA, CDR global scores, and RAVLT. The participants in NC group showed better performance than MCI and AD groups. Here, the MoCA scores seems to be lower than expected which could be presumably due to the linguistic and cultural differences between the original English version and the Chinese version of the scale or the lower education level of Chinese older adults (Yu et al., 2012).

**Table 1.**
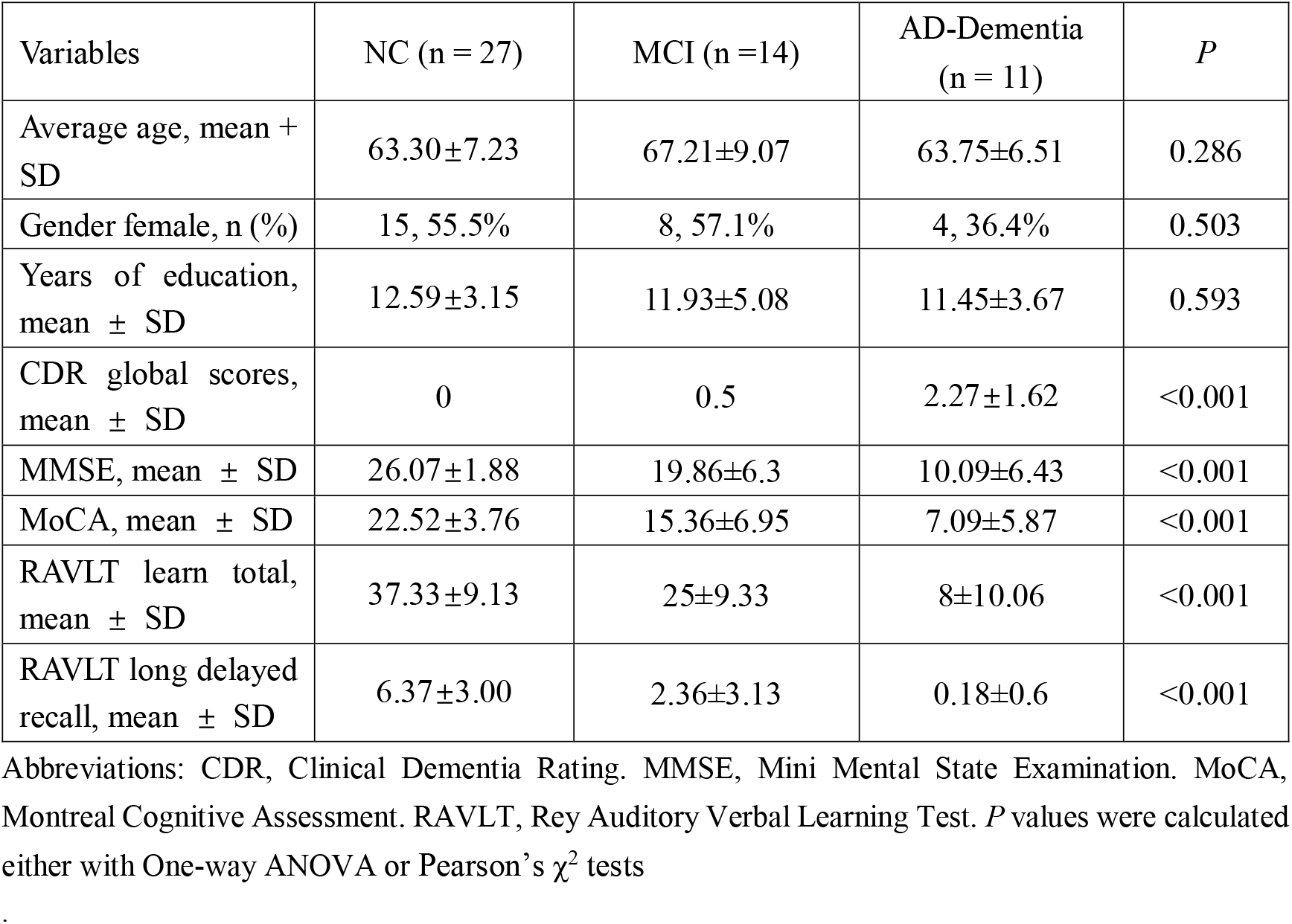
Subject characteristics.

### 3.2. MCI shows increased BBB permeability in hippocampus

**Figure 3** showed VEXI results of hippocampus in MCI and NC groups. Comparing with that in NC group, the apparent diffusion coefficient *ADC’*of hippocampus in MCI showed larger values at each mixing time *t*_m_ and recovered faster as *t*_m_ increased (**Figure 3A&3B**), in which the latter suggests faster water exchange across BBB (*i*.*e*., increased BBB permeability, **Figure 3B**). Further quantitative modeling (**Eq. 1–3**) revealed that the AXR_BBB_ of hippocampus in MCI group (n = 14, group-averaged AXR_BBB_ = 1.63 s^-1^) was significantly larger than those in NC group (n = 27, AXR_BBB_ = 1.21 s^-1^, *p* = 0.042, **Figure 3C**). Besides the hippocampus, no significant BBB changes between NC and MCI group were found in other cortical, subcortical or WM regions.

**Figure 3.**
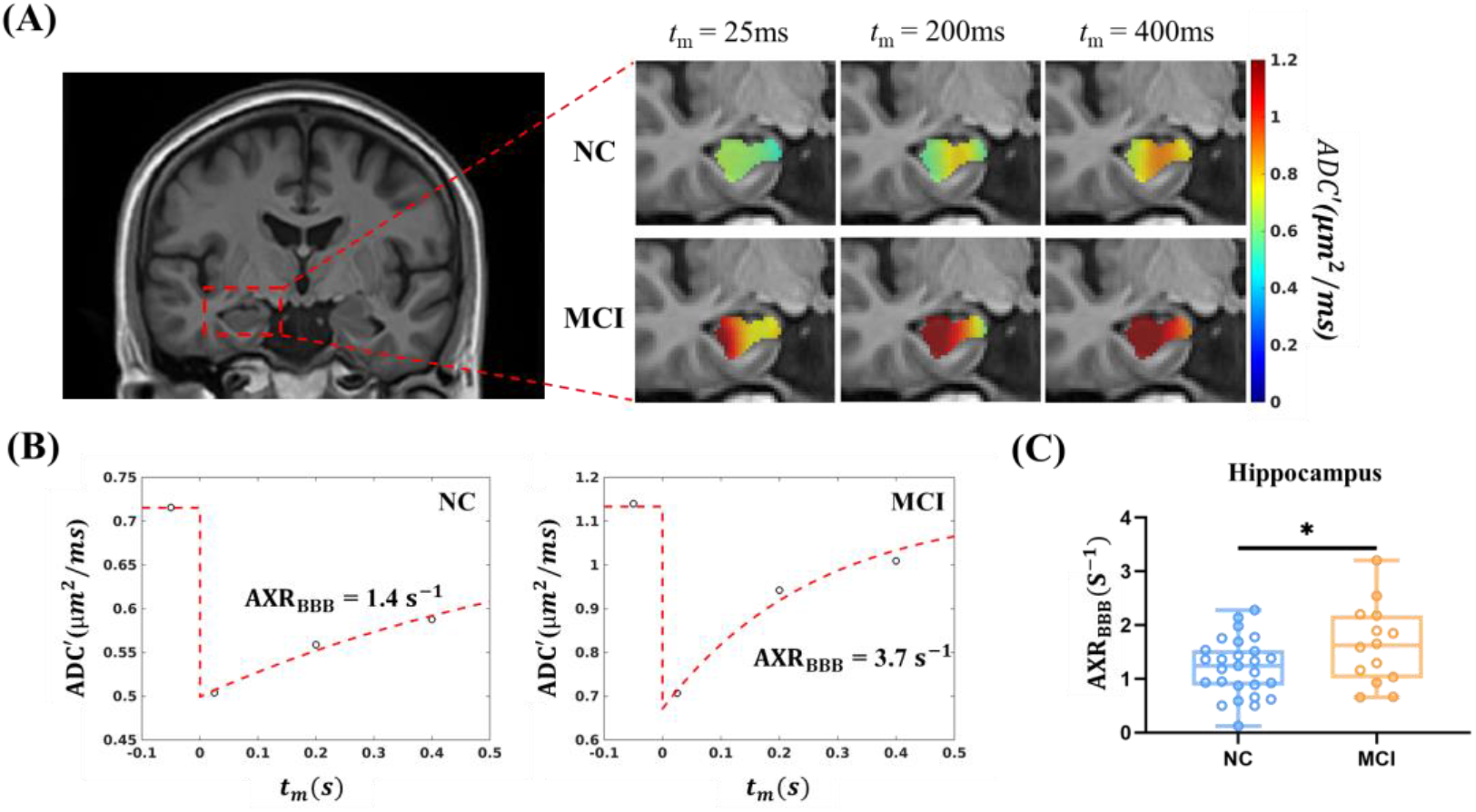
VEXI showed BBB breakdown of hippocampus in MCI group. (A) the averaged ADC’ maps of all subjects in the MCI group and NC group at each mixing time. Here only the results in hippocampus were shown in color. (B) Representative *ADC*′(*t*_*m*_) curves of the hippocampus in the NC (left) and MCI (right) group were shown. Here the first circle at *t*_*m*_ < 0 *s* denoted as the equilibrium state (*b*_f_ = 0 s/mm^2^). (C) Statistical comparison of AXR_BBB_ between NC and MCI in hippocampus. Unpaired Students’ *t*-test. * *p* < 0.05. NC, normal cognition group (n=27); MCI, mild cognitive impairment group (n=14).

### 3.3. The BBB breakdown gets worse and extends to more brain regions from MCI to AD

To investigate the BBB integrity as the disease progresses, we further analyzed the AXR_BBB_ change from NC to MCI, and then to AD. **Figure 4A-C** showed the averaged AXR_BBB_ maps of all subjects in NC, MCI, and AD groups. In hippocampus, we found increased AXR_BBB_ as the disease progressed from MCI to AD (**Figure 4A**), suggesting more damage of BBB with severe cognitive dysfunction. Further ANOVA analyses revealed significant AXR_BBB_ changes in the hippocampus (F = 4.75, *p* = 0.013, **Figure 4D**), which increased by 56.2% from NC to AD (*p* = 0.016, Tukey’s post hoc test). In addition, the BBB permeability showed spatial patterns inside the hippocampus: the subregions close to CA1 showed the earliest AXR_BBB_ changes from NC to MCI. Due to the low spatial resolution of the current VEXI method, we didn’t pursue further quantitative analysis of hippocampal subregions. In addition to hippocampus, thalamus and OFC also showed significantly increasing AXR_BBB_ (by 59.4% and 130.0%, respectively) from NC to AD (*p* = 0.0006 and *p* = 0.0001, respectively, Tukey’s post hoc test after ANOVA analysis) but didn’t show detectable BBB permeability changes from NC to MCI even in the unpaired Students’ t test (**Figure 4E&4F**).

**Figure 4.**
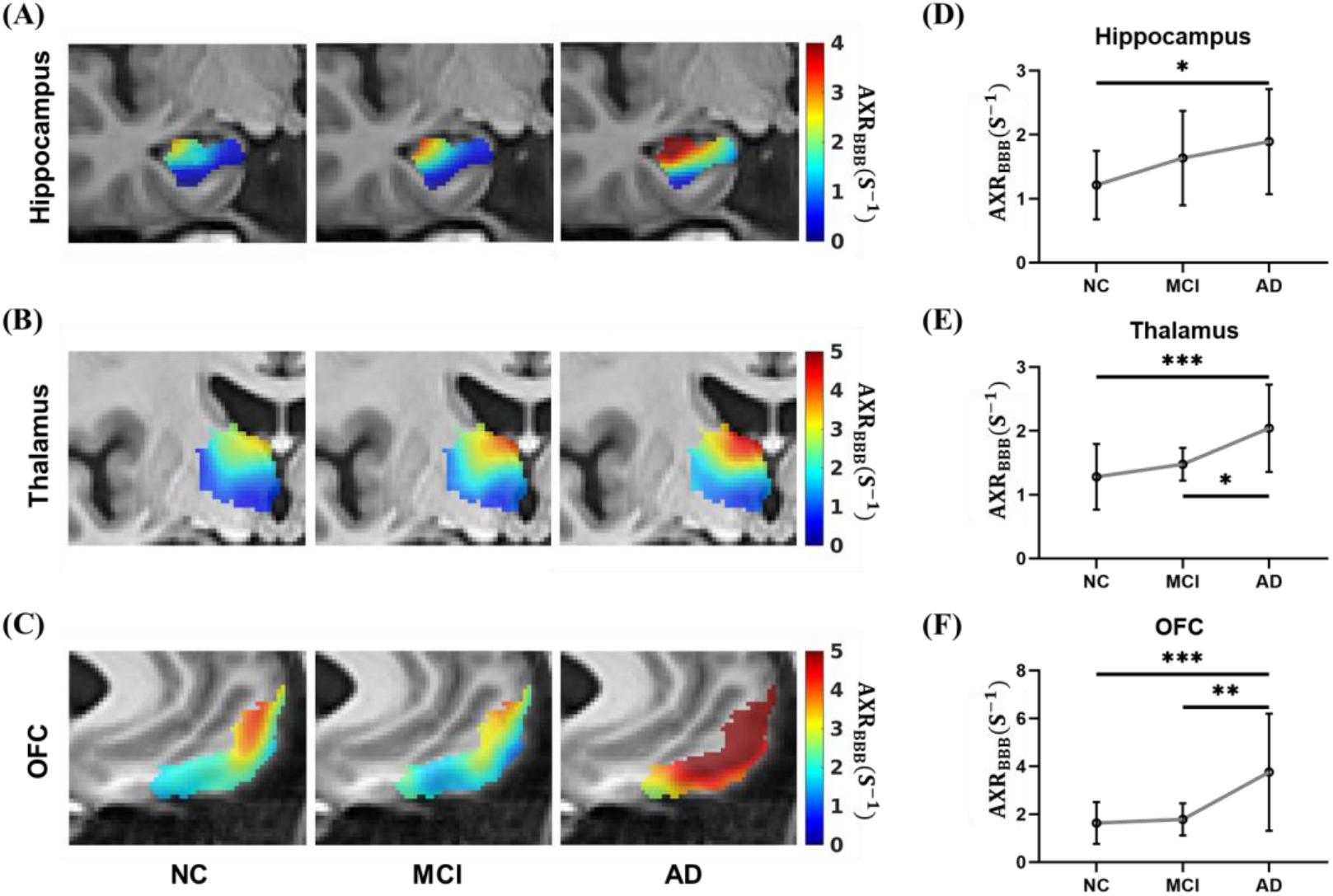
VEXI showed BBB breakdown along with disease progression. (A-C) Averaged AXR_BBB_ maps of all subjects in the NC, MCI and AD groups in hippocampus, thalamus and OFC in the template space. Statistical comparison of AXR_BBB_ values in hippocampus (D), thalamus (E), and OFC (F) among the NC (n=27), MCI (n=14) and AD (n=11) groups. *p*, significance by ANOVA followed by Tukey’s post hoc tests. * *p* < 0.05, ** *p* < 0.01, *** *p* < 0.001. NC, normal cognition group; MCI, mild cognitive impairment group; AD, Alzheimer’s disease.

To eliminate the potential impact of different sex composition between AD and NC (though not significant), we performed permutation test by randomly choosing 7 females out of 15 in the NC group to ensure the NC group has roughly the same sex ratio as the AD group. Among all the 6435 combinations, 100% of them in thalamus and OFC and 94.6% of them in hippocampus showed significantly smaller AXR_BBB_ than the AD group (*p* < 0.05, unpaired Students’ t-test), demonstrating the above results were not biased by the sex composition variance.

Hippocampal atrophy is a well-established biomarker in AD. In this study, we also found the significant reduction of hippocampus volume from NC to MCI and from MCI to AD. In fact, all the selected brain regions showed significant volume reduction in the AD group (**Figure S1**). To further demonstrate that our AXR_BBB_ results in the template space were not affected by the brain morphology changes and these registration steps from subject’s native space to template space, the same ROI-based VEXI parameter analysis were also performed in in the subjects’ native space (see methods). The hippocampus and other brain ROIs in each subject were manually drawn by one experienced neurologist and the CSF voxels were also carefully removed. Further ANOVA test on the three groups found significant differences in AXR_BBB_ only on the same three brain regions (hippocampus, thalamus, and OFC) (Supplementary Information **Figure S2**). These results agree with the findings in the template space (Figure 4) and further demonstrates the reliability of our results in the template space.

### 3.4. Correlation between BBB breakdown and cognitive dysfunction

Linear regressions were used to assess potential correlations between BBB impairment and cognitive dysfunction in all subjects and subjects except AD group. As shown in **Figure 5** although the AXR_BBB_ of all the three brain regions showed significant and negative correlations with the MoCA score in all subjects (*r* = -0.44, *p* = 0.001 in hippocampus, *r* = -0.50, *p* = 0.0002 in thalamus and *r* = -0.53, *p* < 0.0001 in OFC), only in hippocampus this correlation remained significant in the absence of AD group (*r* = -0.32, *p* = 0.043), suggesting that the worse cognitive function is associated with larger hippocampal BBB impairment while the BBB impairment is related to cognitive dysfunction in thalamus and OFC only at a late stage of the disease. Further statistics revealed that the AXR_BBB_ of hippocampus, thalamus, and OFC in all subjects are all significantly associated with cognitive performance, such as MMSE, MoCA, RAVLT learn total and RAVLT long delayed recall, independent of age, sex and education (**Table 2**). We also did the correlation analysis between AXR_BBB_ of the three brain regions and other neurological tests in the absent of AD group. Similar as the above MoCA result, only in hippocampus the correlations remained significant or close to significant (except for RAVLT long delayed recall) (**Table S1**).

**Table 2.**
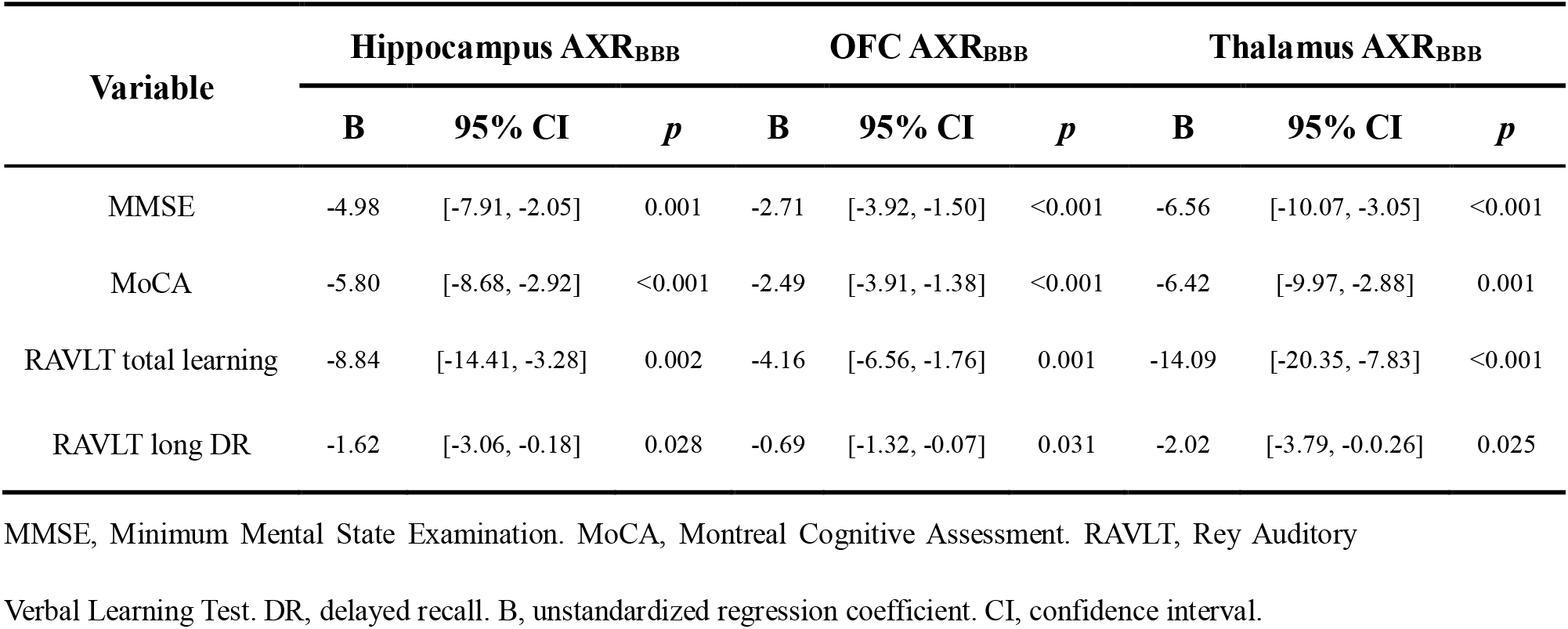
Linear regression of BBB impairment and cognitive performance.

**Figure 5.**
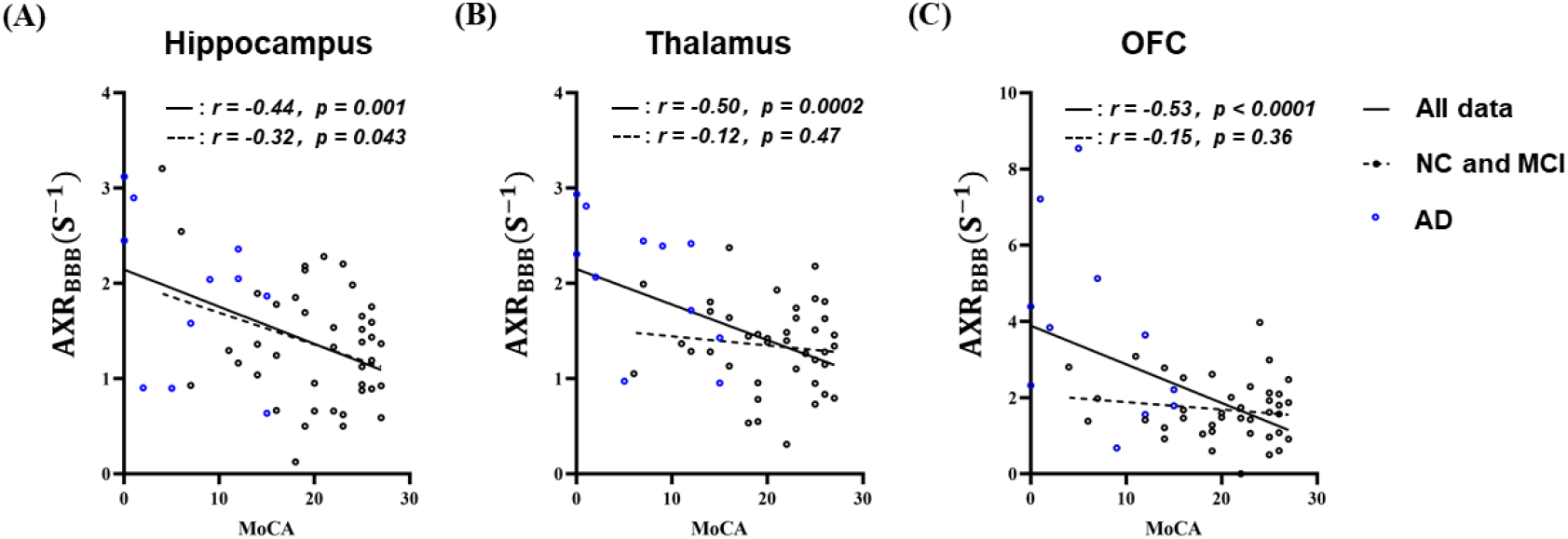
Larger AXR_BBB_ was found to be associated with lower MoCA score in hippocampus(A), thalamus(B) and orbital frontal cortex(C). However, in the absence of AD data, the correlation between AXR_BBB_ and MoCA only remained in hippocampus. Here each dot denoted the data from each subject (black for NC and MCI, blue for AD). The solid and dotted line denoted the linear regression results in all the data and in the absence of AD data, respectively, along with the correlation coefficients (and *p* values) labeled after solid and dotted lines.

### 3.5. Changes in other VEXI metrics along the progression of AD

Significant increase in ADC_0_ was found as the disease progress in hippocampus, thalamus, and OFC (**Figure 6A-6C**). ANOVA analyses revealed significant ADC_0_ changes in the hippocampus (*F* = 14.93, *p* < 0.0001, **Figure 6A**), which increased by 44.2% from NC to AD (*p* < 0.0001, Tukey’s post hoc test). OFC showed the similar patterns from NC to AD with ADC_0_ increased by 16.4% (ANOVA test, *F* =7.37, *p* = 0.0016, **Figure 6C**). In thalamus, significant ADC_0_ increase was found from NC to MCI by 6% though no increase was found from MCI to AD (F = 4.65, *p* = 0.014, **Figure 6B**). All the other 5 brain ROIs, including caudate, striatum, IC, CC, and WSWM, also showed significant ADC_0_ increases from NC to AD (**Figure S3**). As for filter efficiency *σ*, significant decreases in *σ* from NC to AD were only found in thalamus (**Figure 6E**) and caudate nucleus (**Figure S3**). No significant changes in *σ* were found in other brain regions.

**Figure 6.**
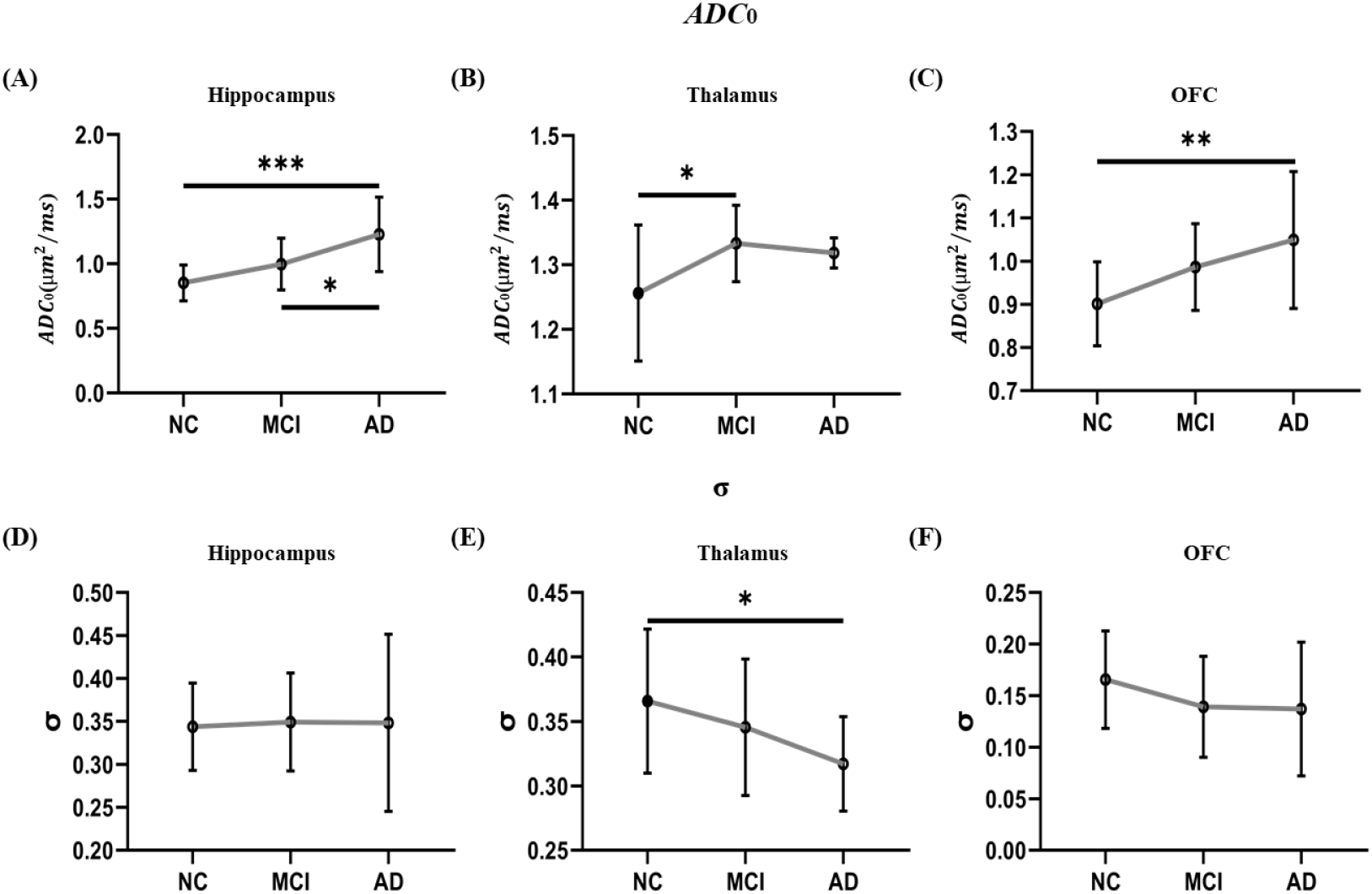
Changes in the other VEXI metrics, ADC_0_ and *σ*, along the progression of AD. Significant ADC_0_ increase was found in hippocampus (A), thalamus (B), and OFC (C) in the progression of AD. No significant *σ* changes were found in hippocampus (D) and OFC (F), meanwhile thalamus (E) shows a significant decrease from NC to AD. *p*, significance by ANOVA followed by Tukey’s post hoc tests. * *p* < 0.05, ** *p* < 0.01, *** *p* < 0.001. NC, normal cognition group (n=27); MCI, mild cognitive impairment group (n=14); AD, Alzheimer’s disease (n=11).

## 4. Discussion

BBB breakdown is an essential pathophysiological process in AD and leads to the accumulation of potentially neurotoxic blood-derived products in the brain that normally do not go across BBB. Most neuroimaging studies that assessing BBB integrity in AD need the administration of a contrast agent to characterize the vascular leakage of the contrast agent. In this study, we first assessed the feasibility of vascular water exchange imaging (VEXI), a special diffusion MRI method assessing the speed of water exchange across BBB without the requirement of contrast agent, in detecting the BBB breakdown in AD. We found VEXI could detect the subtle BBB breakdown in the MCI group in comparison with NC, which specifically happened in the hippocampus. We then found BBB breakdown got worse from MCI to AD and extended to the thalamus and OFC regions. Furthermore, the BBB permeability to water detected by VEXI showed a significant correlation with the cognitive dysfunction. Taken together, our results have demonstrated the feasibility of VEXI in the detection of BBB breakdown in AD and VEXI as a potential contrast-free neuroimaging method in the long-term studies of a large population.

Hippocampus, a region critical for learning and memory, shows BBB breakdown in aging and early AD (Sweeney et al., 2018). It has been demonstrated that BBB breakdown is an early event in aging human brain that begins in the hippocampus using dynamic-contrast enhanced MRI (DCE-MRI) and this BBB breakdown is correlated with injury to BBB associated pericytes and may contribute to cognitive impairment (Montagne et al., 2015). In AD patients, BBB breakdown in hippocampus was detected by using DCE-MRI methods and shown as an early sign of cognitive dysfunction independent of Aβ and/or tau changes (Nation et al., 2019) but related to APOE4 (Montagne et al., 2020). In this study, we have also found faster water exchange across BBB (i.e., AXR_BBB_) in the hippocampus in AD, which demonstrates the BBB breakdown in the hippocampus and agrees with these literatures. More importantly, the BBB breakdown in the hippocampus detected in VEXI correlates with cognitive dysfunction and this correlation remains in the absence of AD data, suggesting its potential value as a biomarker of cognitive impairment even in the early AD phase.

In this study, we have also demonstrated the BBB breakdown of the OFC in AD, which is in line with the previous studies using DCE-MRI methods (van de Haar et al., 2017, 2016). The orbital frontal cortex is a region involved in awareness and metacognitive processes, in line with an abundant literature (McGlynn and Schacter, 1989). Studies with PET (Salmon et al., 2006) and SPECT (Mimura and Yano, 2006) have already reported a link between anosognosia in AD and OFC dysfunction. A previous research has shown a significant loss of pericytes in the frontal cortex in AD patients, which is correlated with BBB leakage (Sengillo et al., 2013).

In the comparison between AD and NC, the thalamus also showed significant BBB breakdown using VEXI, which is not reported in recent DCE-MRI studies (Montagne et al., 2020, 2015; Nation et al., 2019). This is not surprising as a previous study on AD rat model found that water exchange across BBB was a more sensitive biomarker than the vascular leakage of MRI contrast agent (e.g., transfer constant *K*^trans^) in the detection of subtle BBB breakdown (Dickie et al., 2019). Since water molecule is much smaller than contrast agent molecule, the biomarker of water exchange across BBB showed BBB breakdown in several brain regions (including hippocampus, thalamus, and cortex) in this AD rat model while *K*^trans^ failed to detect any BBB breakdown (Dickie et al., 2019). Indeed, some other studies using DCE-MRI have also reported the BBB leakage in deep gray matter and cortex (van de Haar et al., 2017). Thalamus, once viewed as passively receiving information from the basal ganglia limbic system and cerebellum, then relaying information to the cerebral cortex, is becoming increasingly acknowledged as actively regulating the information transmitted to cortical areas (B. et al., 2013). Evidence reveals that lesions to higher-order thalamic areas, such as the pulvinar and mediodorsal nucleus, can produce severe attention and memory deficits (Baxter, 2013; Bradfield et al., 2013), suggesting an important role for the thalamus in cognition. Recent studies have also shown the atrophy of thalamus in AD, which has also been proven in this study, and its potential role in the cognitive dysfunction (De Jong et al., 2008; Pardilla-Delgado et al., 2021).

For the other VEXI metrics, *ADC*_0_ shows significant increases along the progression of AD in all the selected brain regions, which agrees with the observed mean diffusivity (MD) or apparent diffusion coefficient (ADC) in MCI (or AD) in comparison with NC in previous studies (Altamura et al., 2016; Kantarci et al., 2005; Scola et al., 2010; Takahashi et al., 2017). Hippocampal ADC has been demonstrated as an effective predictor of the conversion from amnestic MCI to AD(Kantarci et al., 2005). Beyond hippocampus, a further study has also found the MD increase in the whole brain and some manually segmented ROIs along the trajectory from NC to MCI and to AD (Scola et al., 2010). The filter efficiency *σ* is a more complex parameter, which can be affected by the vascular water mole fraction and the ADCs of water in both intra- and extra-vascular space (see Eq. 5 in Bai et al., 2020). Both global and regional cerebra hypoperfusion has been demonstrated in AD and during the preclinical phase of AD (i.e., MCI) (Austin et al., 2011; Binnewijzend et al., 2016).Hypoperfusion could result to decreased ADC of intravascular water and/or decreased vascular water mole fraction (Le Bihan, 2019; Zhu et al., 2020). These factors, along with the increased ADC of extravascular water could explain the decreased *σ* in some brain regions from NC to AD, though most regions don’t show significant changes. However, conflict results have also been reported. In a recent study using IVIM-DWI method, the detected vascular water mole fraction was found to increase with cognitive decline (Bergamino et al., 2020). The good news is that these two VEXI metrics are independent from AXR_BBB_ estimation (Eq. 4) and the changes in *ADC*_0_ or *σ* will not affect the accuracy of AXR_BBB_ estimation.

Several limitations and future works of this study should be clarified. At first, though previous study has demonstrated the increased BBB permeability to water is associated with the reduced expression of the tight junction protein occludin in AD rat model (Dickie et al., 2019) and CSF Aβ42 concentration levels in healthy older human adults (Gold et al., 2021), it is still highly desired to provide further pathophysiological explanations of the increased BBB permeability to water molecule in human AD. For instance, CSF levels of soluble platelet-derived growth factor receptor β (sPDGFRβ) are one of the markers to assess the BBB-associated pericytes that play a key role in maintaining the BBB integrity (Sagare et al., 2015) and can be used to compare with the VEXI results in future. Another limitation is the small sample size, though careful statistics were performed. It is highly desired to test VEXI in a large sample and in longitudinal studies. Third, AXR_BBB_ could potentially be biased from intravascular water efflux rate constant *k*_fs_ by the difference in vessel density among different groups (Eq. 4), but this bias (<5%, Bai et al., 2020) is negligible in comparison with the relatively large AXR_BBB_ changes among different groups (>30%). Forth, the current VEXI still suffers from low spatial resolution, which limits further analysis on the subregions of hippocampus, thalamus, and other brain regions. Improving the spatial resolution of VEXI without scarifying the signal-to-noise quality is warranted in future studies. At last, it is interesting to study the age-dependence of the BBB permeability to water molecules using the same MRI method (e.g., VEXI) in future. Due to the acquisition parameter differences between this study and our previous work on young and healthy subjects (Bai et al., 2020), it is hard to compare the results in these two studies directly.

## Acknowledgments

We would like to thank all the subjects for their participation in the study. We appreciate the help and work of all related clinicians, statisticians, imaging technicians and coordinators who participated in this study. This study is supported by grants from National Natural Science Foundation of China (NSFC) (Grant No. 81873894, Grant No. 82111530201), Beijing Municipal Science & Technology Commission (Grant No. Z181100001518005), Natural Science Foundation of Zhejiang Province, China (Grant No. LR20H180001), Strategic Priority Research Program of the Chinese Academy of Sciences (XDB39000000), and the MOE Frontier Science Center for Brain Science & Brain-Machine Integration, Zhejiang University.

