## Supplementary Information for "Vascular-water-exchange MRI (VEXI) enables the detection of subtle BBB breakdown in Alzheimer’s disease without MRI contrast agent"

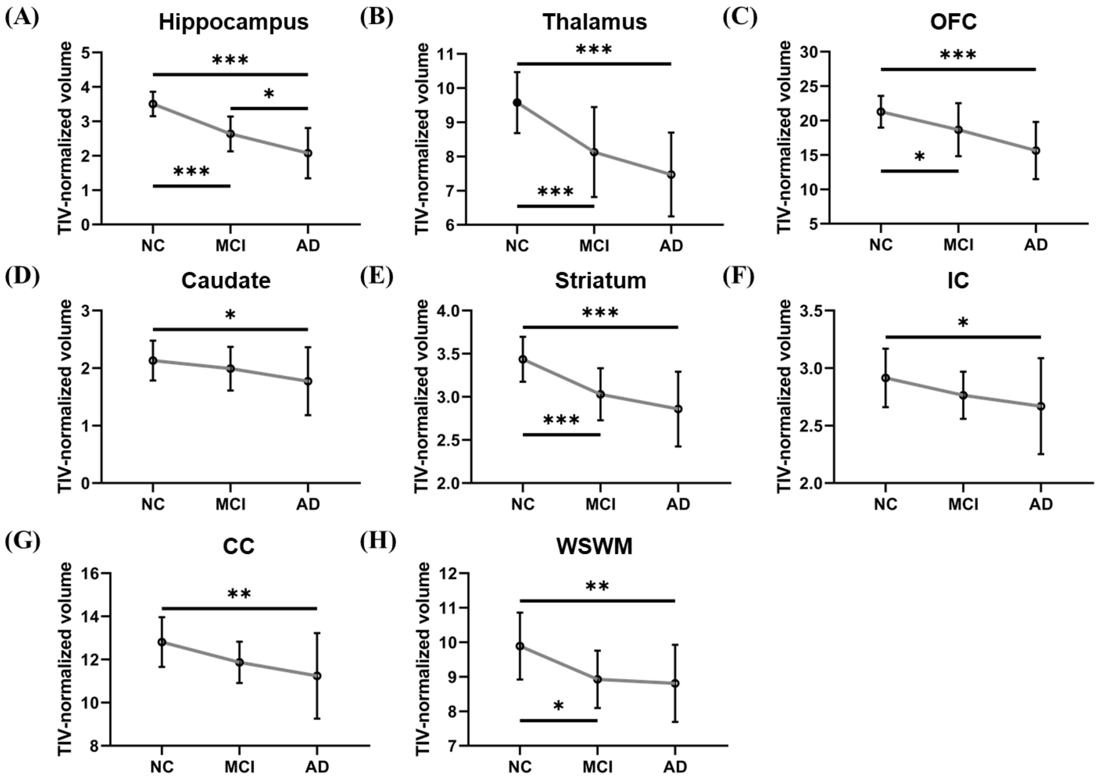


**Figure S1. Brain Atrophy in the MCI and AD groups.** Relative to NC, MCI and AD groups exhibited significantly smaller brain volume in hippocampus (A), thalamus (B), OFC(C), striatum (E) and WSWM (H). As for caudate (D), IC (F) and CC (G), significant volumetric decreases were only found in the AD group. TIV-Normalized volumes were subsequently multiplied by a factor of $10^3$ to facilitate ease of comparisons. $p$, significance by ANOVA followed by Tukey's post hoc tests. * $p < 0.05$, ** $p < 0.01$, *** $p < 0.001$. NC, normal cognition group (n=27); MCI, mild cognitive impairment group (n=14); AD, Alzheimer's disease (n=11). TIV, total intracranial volume. OFC, orbital frontal cortex. IC, internal capsule. CC, corpus callosum. WSWM, watershed white matter fibers.

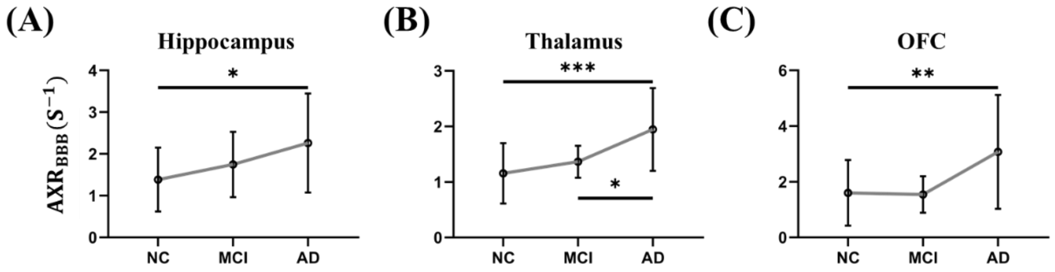


**Figure S2.** In the subjects' native space, VEXI also showed BBB breakdown along the AD progression only in the three brain regions (hippocampus, thalamus, and OFC). Statistical comparison of $AXR_{BBB}$ values in hippocampus (A), thalamus (B), and OFC (C) among the NC, MCI, and AD groups. *p*, significance by ANOVA followed by Tukey's post hoc tests. * $p < 0.05$, ** $p < 0.01$, *** $p < 0.001$. NC, normal cognition group (n=27); MCI, mild cognitive impairment group (n=14); AD, Alzheimer's disease (n=11).

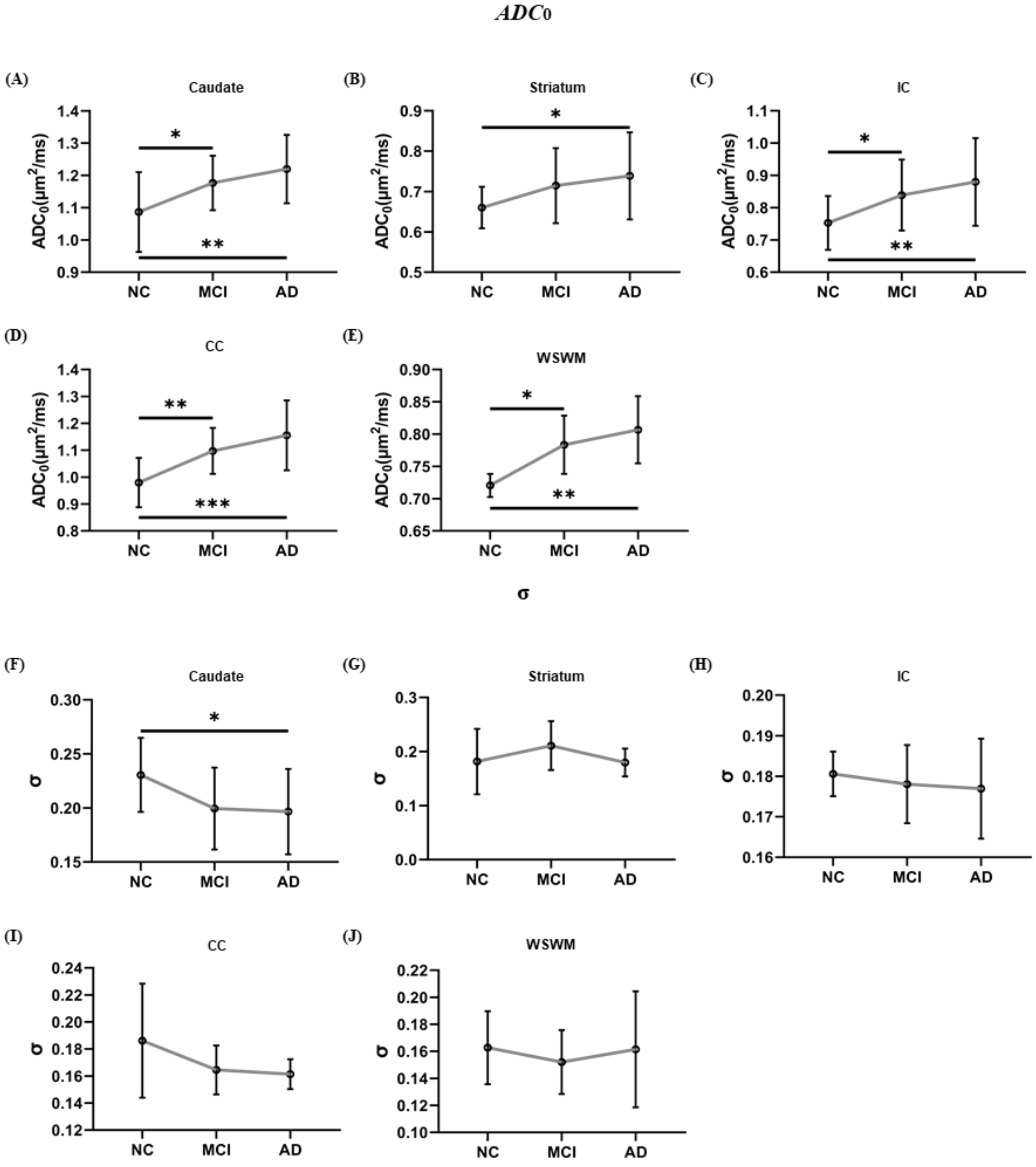


**Figure S3.** Regional $ADC_0$ and σ changes from NC to MCI and AD. (A-E) significant $ADC_0$ increase was found in selected ROIs including caudate, striatum, IC, CC and WSWM. However, no significant σ change was found in these brain ROIs (G-J), except for caudate(F). *p*, significance by ANOVA followed by Tukey's post hoc tests. * $p < 0.05$, ** $p < 0.01$, *** $p < 0.001$. NC, normal cognition group (n=27); MCI, mild cognitive impairment group (n=14); AD, Alzheimer's disease (n=11). OFC, orbital frontal cortex. IC, internal capsule. CC, corpus callosum. WSWM, watershed white matter fibers.

| Variable | Hippocampus $AXR_{BBB}$ | | | OFC $AXR_{BBB}$ | | | Thalamus $AXR_{BBB}$ | | |
|---|---|---|---|---|---|---|---|---|---|
| | **B** | **95% CI** | ***p*** | **B** | **95% CI** | ***p*** | **B** | **95% CI** | ***p*** |
| MMSE | -3.34 | [-5.87, -0.80] | 0.011 | -0.92 | [-2.98, 1.14] | 0.37 | 0.54 | [-2.73, 3.80] | 0.74 |
| MoCA | -3.95 | [-7.02, -0.87] | 0.013 | -1.12 | [-3.62, -1.37] | 0.37 | 1.10 | [-2.96, 5.15] | 0.59 |
| RAVLT total learning | -5.40 | [-10.88, -0.08] | 0.053 | -0.33 | [-4.60, 3.95] | 0.88 | -1.81 | [-9.12, 5.50] | 0.62 |
| RAVLT long DR | -1.02 | [-2.87, 0.83] | 0.27 | -0.09 | [-1.29, 1.48] | 0.89 | 0.18 | [-2.47, 2.82] | 0.89 |

**Table S1. Linear regression of BBB impairment and cognitive performance in the NC and MCI groups**. Age, sex and education were controlled for the analysis. MMSE, Minimum Mental State Examination. MoCA, Montreal Cognitive Assessment. RAVLT, Rey Auditory Verbal Learning Test. DR, delayed recall. B, unstandardized regression coefficient. CI, confidence interval.
